# Multiscale Symbolic Morpho-Barcoding Reveals Region-Specific and Scale-Dependent Neuronal Organization

**DOI:** 10.64898/2026.02.28.708657

**Authors:** Sujun Zhao, Yingxin Li, Yufeng Liu, Hanchuan Peng

## Abstract

Neuronal morphology is a central determinant of circuit organization, yet its multiscale complexity has hindered systematic, brain-wide analysis and integration with anatomical context. Here we introduce **Multiscale Morpho-Barcoding (MMB)**, a framework that encodes whole-brain neuronal morphology into symbolic representations spanning cellular geometry, axonal tract routing, arbor organization, and predicted synaptic distributions. Applying MMB to 1,876 fully reconstructed mouse neurons, comprising 3,776 arbors and 2.63 million predicted presynaptic sites, we identify distinct multiscale morpho-patterns that reveal region-specific and scale-dependent principles of neuronal organization across the brain. MMB robustly discriminates major anatomical divisions and resolves canonical thalamic circuit classes beyond what can be achieved using projection strength alone. By transforming complex neuronal geometry into interpretable multiscale representations, MMB provides a general framework for systematic comparison of neuronal structure and for integrating morphology with connectivity and function at whole-brain scale.

## Introduction

Neuronal morphology plays a fundamental role in shaping circuit computation, influencing how neurons integrate inputs, route signals, and establish long-range connectivity [1,2]. Advances in whole-brain imaging and automated reconstruction have enabled the acquisition of thousands of single-neuron morphologies across the mammalian brain [3-15]. However, the resulting data have exposed a key limitation in current analysis approaches: there is no general representation that captures neuronal structure across hierarchical scales while enabling systematic, brain-wide comparison and integration with anatomical context [16,17,18].

Existing morphometric analyses typically focus on predefined feature sets or individual structural scales, such as global geometry [19,20], local branching statistics [21], or synapse distributions [22, 23]. While effective for specific questions, these approaches struggle to capture how morphological organization at different scales jointly contributes to neuronal identity and regional specialization. As a result, it remains unclear how structural diversity is distributed across the brain, how it varies across scales, and how it relates to large-scale anatomical organization [16, 24].

Addressing this challenge requires a representation that is both multiscale and interpretable, enabling neurons to be compared across regions without collapsing their hierarchical structure into a single descriptor. Such a representation should preserve scale-specific information while supporting quantitative, brain-wide analysis.

Here we introduce Multiscale Morpho-Barcoding (MMB), a framework that encodes neuronal morphology into symbolic barcodes spanning four hierarchical levels: whole-cell geometry, axonal tract routing, arbor organization, and predicted presynaptic distributions. By applying MMB to a large set of fully reconstructed mouse neurons, we perform a systematic, whole-brain analysis of multiscale morphological organization. Our results reveal scale-dependent and region-specific structural patterns, demonstrating that neuronal morphology is organized along multiple partially independent dimensions across the brain.

## Result

### Construction of a multiscale neuronal morphology framework

We assembled and standardized a large-scale light microscopy dataset of whole mouse brains, comprising imaging data from more than 200 specimens and approximately 3.7 petavoxels in uncompressed form [25]. Neurons were sparsely labeled across multiple regions, including cortex (CTX), thalamus (TH), cerebral nuclei (CNU), and striatum (STR). Whole-brain neuronal morphologies were reconstructed semi-automatically using the cloud-based Collaborative Augmented Reconstruction (CAR) platform [26], which supports multiscale morphometric extraction from whole-brain anatomy to synaptic resolution through iterative reconstruction and validation [16, 27]. To enable consistent spatial comparison, all reconstructions were registered to the Allen Common Coordinate Framework v3 (CCFv3) [28] using the cross-modality registration tool mBrainAligner [29,30].

From this pipeline, we obtained a curated dataset of 1,876 neurons with complete morphologies and associated hierarchical subcellular structures, hereafter referred to as **SEU-A1876** (**Fig. 1A**). These neurons are predominantly located in TH (37.2%), CTX (24.1%), and caudoputamen (CP; 16.8%), providing broad coverage of major forebrain regions.

**Fig 1.**
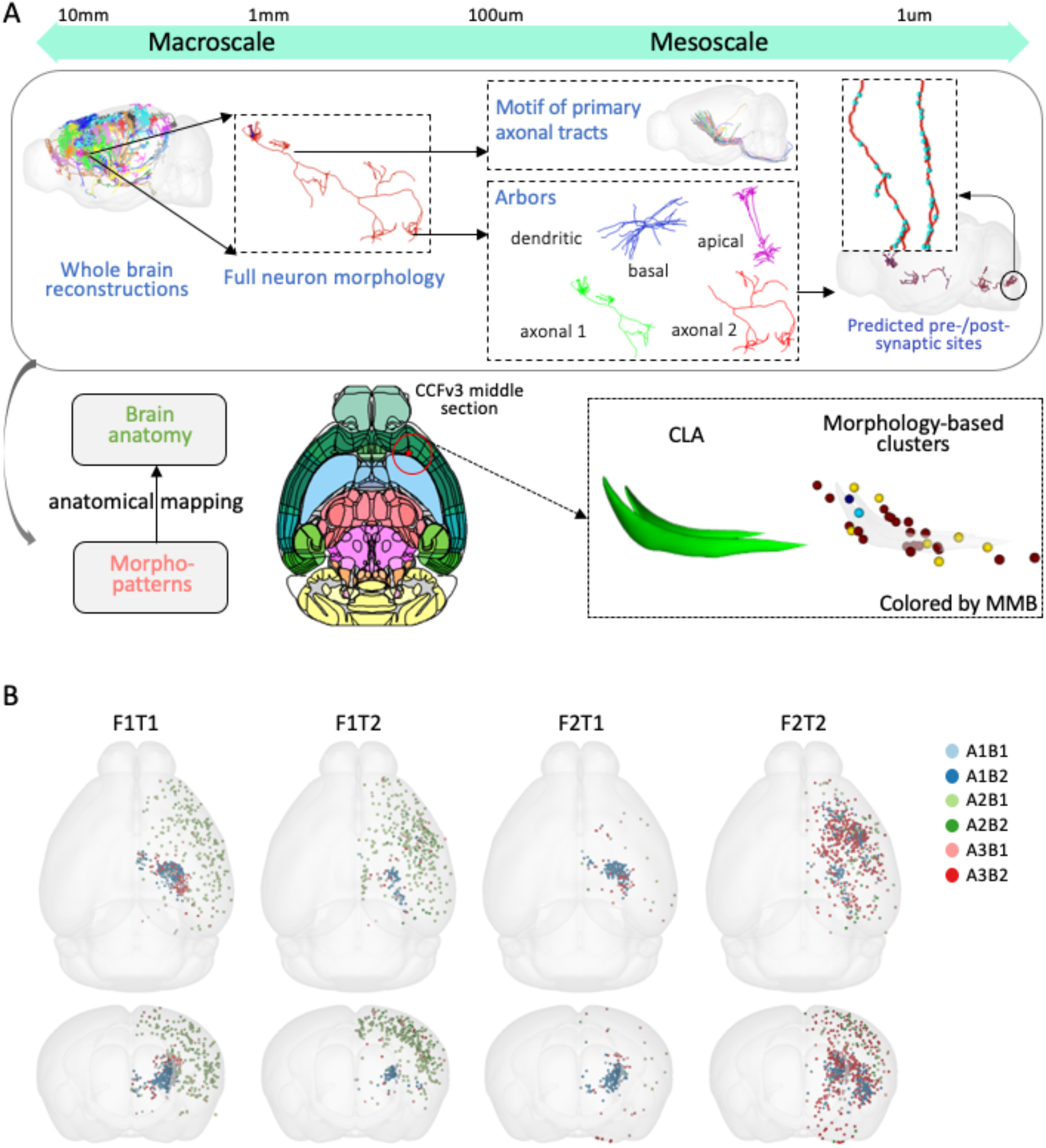
Scheme of Multiscale-Morpho-Barcoding of neuronal patterns and its intended usage to detect associations with other data modalities. **A**. Upper panel: a hierarchical view of neuronal morphology across multiple scales. The complete whole-brain reconstruction includes full morphology, primary tracts, arbors (apical, basal, and axonal domains), and predicted presynaptic sites, spanning from macroscopic to mesoscopic levels. Lower panel: the goal of cross-scale studies to establish correspondences between morphology and brain anatomy, using claustral (CLA) neurons as an example. **B**. Distribution of 24 MMBs across brain regions at the whole brain level, visualized by soma locations. Six elaborated structural modules (A#B#) are embedded within 4 global structural modules (F#T#).

Within this framework, we analyzed neuronal morphology across four hierarchical structural levels: whole-cell morphology, primary axonal tracts, subcellular arborization (dendritic and axonal), and predicted presynaptic sites (**Fig. 1A**). For SEU-A1876, this yielded 1,876 full morphologies, 1,876 primary axonal tracts, 1,876 dendritic arbors, 3,776 axonal arbors, and 2.63 million predicted presynaptic sites [16]. These representations enable systematic characterization of anatomical organization across scales, allowing both conserved (stereotyped) and variable (diverse) morphological patterns to be examined at whole-brain resolution.

For each structural level, we extracted scale-appropriate feature vectors comprising 21 dimensions for full morphology, 6 for primary axonal tracts, 54 for arbors, and 7 for predicted presynaptic sites (**Methods**). Together, these features support hierarchical classification across structural levels, forming the basis of multiscale morpho-barcoding (MMB). This representation preserves scale-specific information while enabling integrated analysis, and effectively captures the spatial and anatomical preferences of neurons across the brain (**Fig. 1B**).

### Hierarchical morpho-organization symbolized and quantified by Multiscale Morpho-Barcoding

To identify the most informative morphological descriptors at each hierarchical scale, we selected the top three Minimum Redundancy–Maximum Relevance (mRMR) features [31] and mapped them onto the Allen CCFv3 template. For each scale, this produced a three-dimensional, brain-wide RGB feature map in which each color channel corresponds to one feature (**Fig. 2A**). These maps reveal spatial gradients and regional enrichment patterns across the brain, providing an intuitive visualization of multiscale morphological organization.

**Fig 2.**
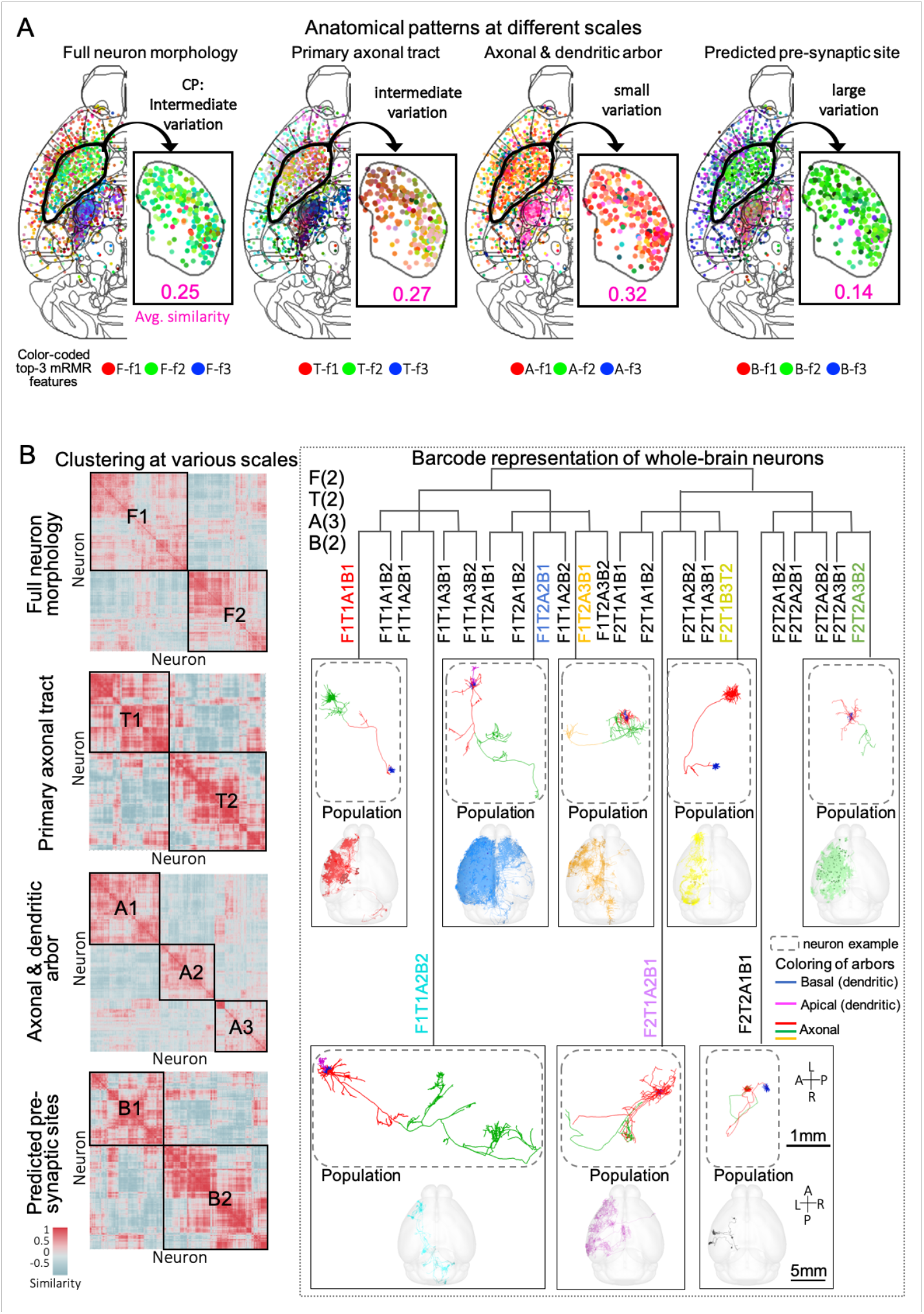
Generation of panoramic multi-barcoding. **A**. Anatomical patterns across four scales: full morphology, primary axonal tract, arbor, and predicted presynaptic site. The three most discriminating features of each scale are shown as colored points on the middle axial section of the CCFv3 atlas. For full morphology, the three features shown are average fragmentation, average remote bifurcation angle, and Hausdorff dimension. For the primary axonal tract, the three features shown are orientation along the anterior–posterior (AP) axis, orientation along the left–right (LR) axis, and Euclidean distance. For the arbor, the three features shown are the number of basal hubs, branch number of axonal arbor 1 (see **Methods** for definition of axonal arbor 1), and distance from the soma to axonal arbor 1. For the predicted presynaptic site, the three features shown are geodesic distance, TEB (terminaux-to-bouton) ratio, and the number of projection target regions. Enlarged insets show feature distributions in the CP region, with variation highlighted in red. **B**. Left panel: clustering results at each morphological scale. Right panel: the hierarchical barcoding architecture for cross-scale morphologies, with representative examples of eight neuron barcodes and their corresponding population visualizations.

At each structural level, neurons were clustered based on their corresponding feature sets (**Fig. 2B**), yielding two clusters for full morphology (F1, F2), two for primary axonal tracts (T1, T2), three for arbors (A1, A2, A3), and two for predicted presynaptic sites (B1, B2). We then encoded each neuron’s multiscale morphological signature as a barcode of the form F#T#A#B#, where each symbol denotes the cluster assignment at a given scale (for example, F2T2A1B2). This representation compactly captures hierarchical anatomical features across multiple scales. In total, this analysis identified 24 distinct neuronal populations, each defined by a unique multiscale morpho-barcode (**Fig. 2B**).

Across the four scales, morphological features exhibited spatially organized and region-specific patterns, with the most pronounced transitions observed at the primary axonal tract level. Compared with the arbor scale, full morphology, tract routing, and presynaptic features showed sharper anatomical boundaries that aligned closely with canonical CCFv3 regions. For instance, a clear boundary between the thalamus (TH) and caudoputamen (CP) was consistently observed across all scales, whereas the cortex–CP boundary was well resolved at all levels except the arbor scale (**Fig. 2A**). Cortex (CTX) displayed greater morphological diversity than other regions whose average feature variance is 12.5 over 10.6 (TH) and 10.5 (CNU), with the strongest spatial gradients emerging at the arbor level (**Table 1**). Within CP, morphological variation was maximal at the presynaptic site level (average similarity across 4 levels: sim = 0.14) and minimal at the arbor level (sim = 0.32), highlighting the scale-dependent nature of neuronal heterogeneity.

To further understand the structural features represented by morpho-barcodes, we can indeed describe each barcode separately across four morpho-scales. At the **full-morphology** scale, the **F module** primarily captures global neuronal size. **F1** neurons are generally larger, exhibiting greater height, width, and depth (**Fig. 3A,B**). In contrast, **F2** neurons are more compact but show higher branch order and larger bifurcation angles, consistent with deeper and potentially more complex branching (**Fig. 3B**). Thus, F clusters reflect not only overall size, but also differences in arborization complexity and spatial expansion.

**Fig 3.**
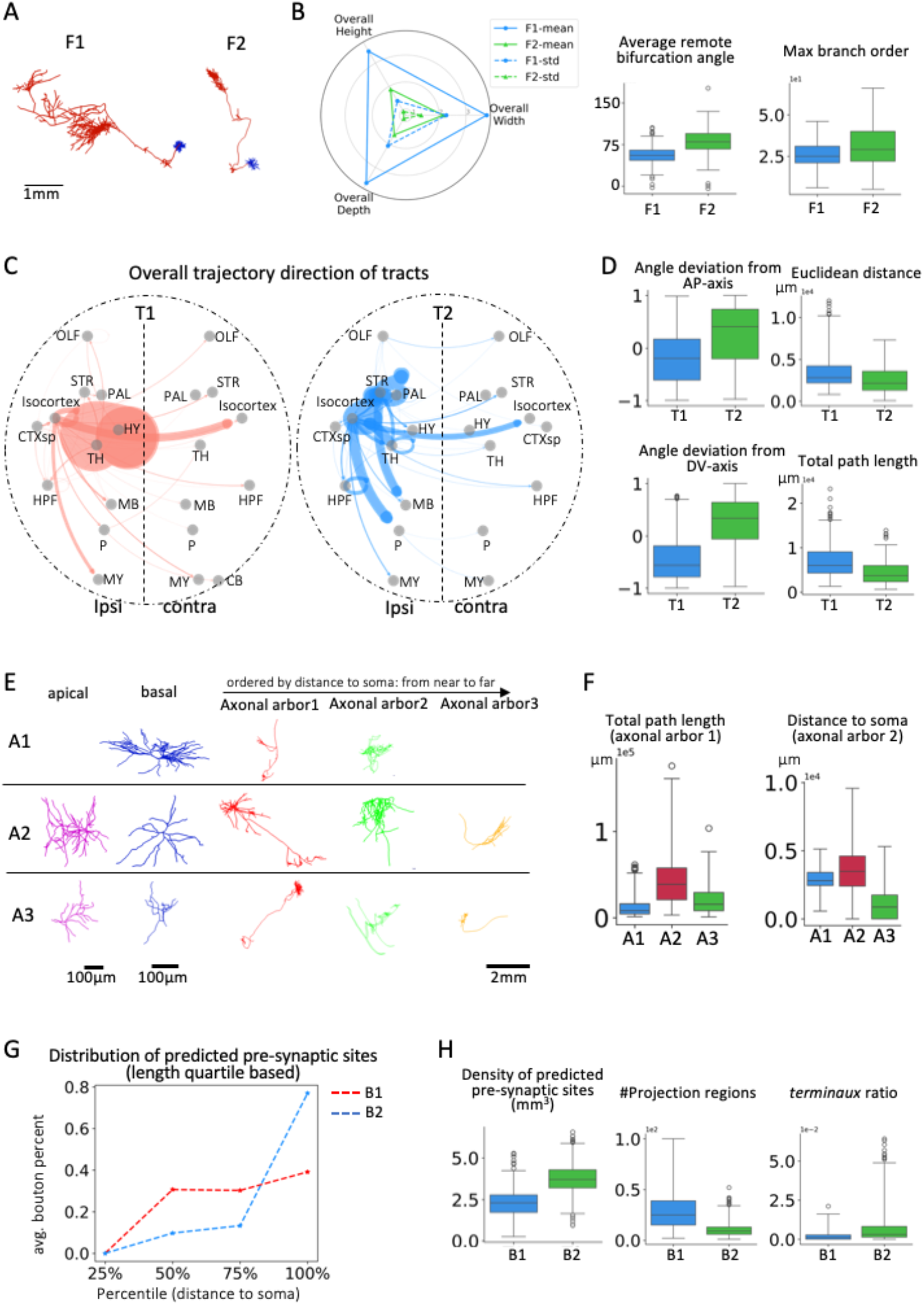
MMB decodes structural features of neuronal morphology. **A**. Examples of F1and F2 type. **B**. Comparison of key morphological features between F1 and F2 neurons. The radar plot on the left summarizes the mean and standard deviation values of neuronal sizes (height, width, depth). Boxplots on the right compare average remote bifurcation angle and max branch order. **C**. Overall projection trajectory directions of T1 and T2 neurons, represented by arrows linking the original and terminal regions of primary axonal tracts; arrow thickness indicates the number of neurons contributing to each trajectory. **D**. Comparison of four tract-scale features: angle deviation from both Anterior-Posterior (AP) and Dorsal-Ventral (DV) axes, Euclidean distance and total path length of tracts. **E**. illustration of apical dendrites, basal dendrites and axonal arbors in three arbor-based clusters (A1-A3). **F**. Comparison of representative arbor-level features across clusters, including total path length of axonal arbor 1, distance from soma to axonal arbor 2. **G**. Spatial distribution of predicted pre-synaptic sites quantified by soma-distance quartiles, shown as the average percentage of sites within specific distance range. **H**. Comparison of key features between B1 and B2 neurons: density of predicted pre-synaptic sites, total number of projection regions, and terminaux ratio. The features above are selected by mRMR.

At the **primary-tract** scale, the **T module** captures long-range routing and projection breadth. **T1** neurons preferentially target thalamic and hypothalamic regions, whereas **T2** neurons exhibit more distributed projection patterns involving striatum, pallidum, isocortex, midbrain, pons, and additional regions (**Fig. 3C**). Consistently, **T2** neurons show larger angular deviations from the anterior–posterior and dorsal– ventral axes, indicating broader trajectory dispersion (**Fig. 3D**). Notably, **T1** neurons tend to have longer primary axonal tracts, suggesting long-distance routing despite more restricted target sets, whereas **T2** neurons project to a wider set of regions (**Fig. 3C,D**).

At the **arbor** scale, the **A module** distinguishes neurons by the organization of apical dendrites, basal dendrites, and axonal arbors. The principal separations are driven by the presence of apical dendrites and the development of **axonal arbor 3** (likely corresponding to the most distal arborization; **Fig. 3E**). **A1** neurons lack apical dendrites and axonal arbor 3, consistent with a relatively small morphological type. By contrast, **A2** neurons exhibit the largest proximal axonal arbors, with **axonal arbor 2** extending the farthest (**Fig. 3F**). In **A3** neurons, axonal arbor 2 lies closest to the soma, suggesting either a smaller overall morphology or prominent branching emerging mid-trajectory along the primary tract (**Fig. 3F**).

Finally, at the **predicted presynaptic-site** scale, the **B module** captures bouton distribution patterns and target diversity. In **B1** neurons, predicted presynaptic sites are distributed across the middle-to-distal portions of the axon and are associated with multiple target regions. In contrast, **B2** neurons show presynaptic sites concentrated near axon terminals, forming spatially clustered bouton distributions that typically correspond to fewer target regions (**Fig. 3G, H**).

Our results reveal a multiscale, symbolic, and quantitative logic of brain organization, in which large-scale anatomical divisions impose global constraints on neuronal morphology, while finer-scale features capture region-specific variability. This MMB representation is not previously available. Indeed, it allows us to use a unified representation to study both stereotyped and diverse aspects of neuronal structure across the brain.

### Neuron organization is region-specific and scale-dependent across the brain

We applied MMB to systematically profile structural organization across brain regions and identified 24 distinct barcodes that represent recurrent configurations of multiscale morphometric modules. Across major brain divisions, neurons exhibit strongly region-specific barcode distributions (**Fig. 4A**). Cortical (CTX) and striatal (CNU) neurons display clearly separable morpho-patterns: CTX neurons are dominated by the F1T1A2B1 and F1T2A2B1 modules, whereas CNU neurons predominantly map to the F2T2A1B2 and F2T2A3B2 modules. In contrast, thalamic (TH) neurons show greater heterogeneity, organized into four principal barcode modules. Specifically, F1T1A1B2 is enriched in the ventral posteromedial (VPM), ventral posterolateral (VPL), medial geniculate (MG), lateral posterior (LP), and submedial (SMT) nuclei; F2T1A1B2 is preferentially observed in VPM, VPL, and SMT; F1T1A3B2 is primarily associated with the dorsal lateral geniculate (LGd), lateral dorsal (LD), and ventral medial (VM) nuclei; and F2T1A3B2 is largely confined to LGd (**Fig. 4B**). Together, these results indicate that neurons within the same anatomical group converge on shared morphological programs, consistent with strong constraints imposed by circuit-level functional demands.

**Fig 4.**
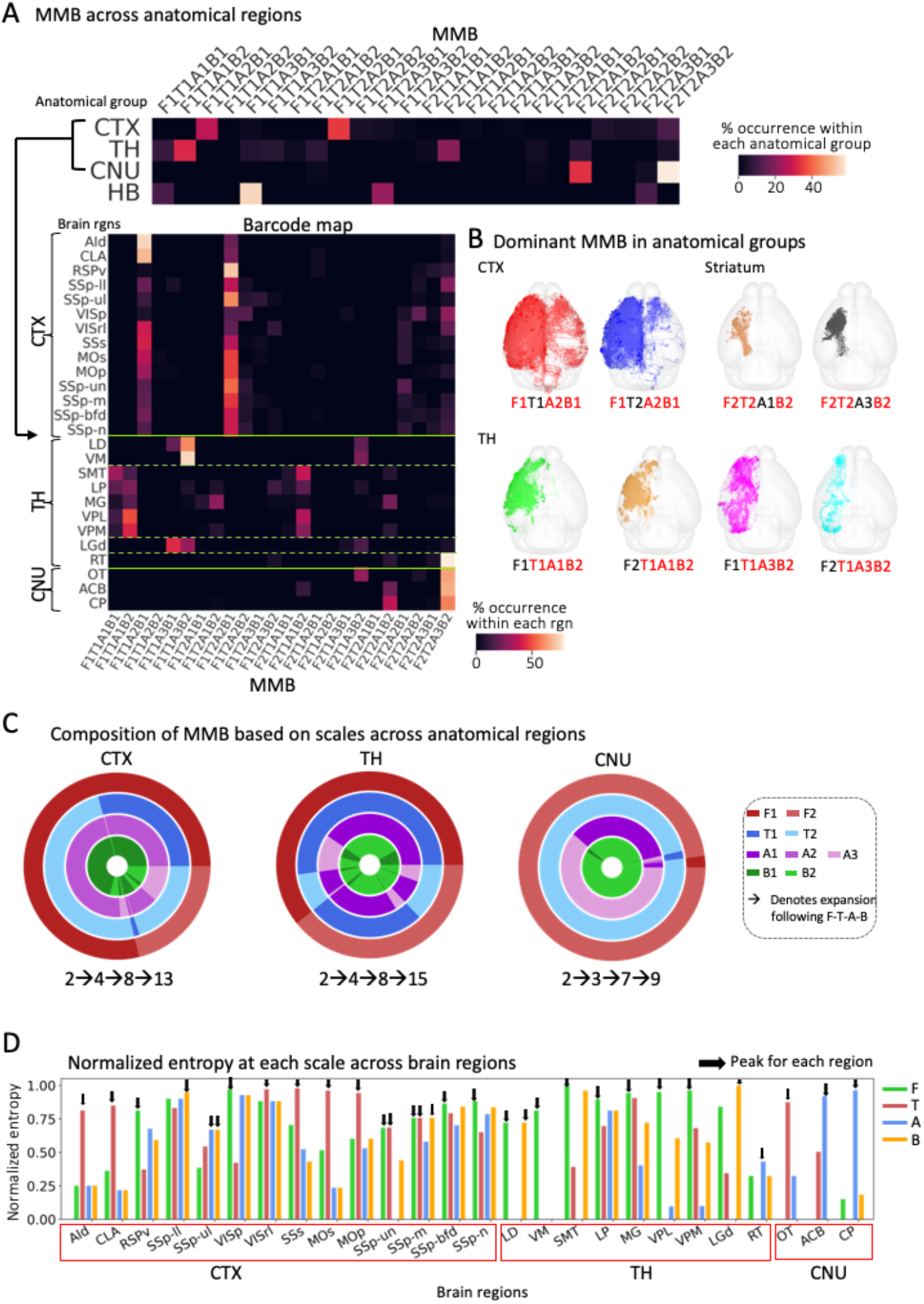
Multiscale morpho-barcoding reveals region-specific neuronal. **A**. Heatmaps show occurrence of morpho-patterns across 3 functional groups and 26 major brain regions (regions with >10 neurons). **B**. Dominant morpho-patterns in 3 functional groups: CTX, STR and TH. **C**. The circled representations show the composition of MMB within major anatomical regions, highlighting the progressive expansion of diversity from full morphology level down to predicted pre-synaptic site level. Taking CTX as an example, the number of barcodes has increased sequentially from 2 to 4, then to 8, and finally to 13, following the progression from full morphology, primary axonal tract, arbor, and predicted pre-synaptic site, indicating an expansion ratio from 2 to 1.625. **D**. Normalized entropy at single morphological scale level for each brain region is shown, with arrows indicating the scale with maximal normalized entropy within each brain region.

To localize the scales at which these region-specific differences emerge, we tracked cluster expansion from the full morphology scale down to predicted presynaptic-site scale, spanning cellular to subcellular organization (**Fig. 4C**). Normalized entropy at each scale is computed as a quantitative measure of diversification (**Table 2, Method**). Bootstrap resampling across most brain regions shows normalized entropy estimates are stable, with narrow confidence intervals and small overlaps across scales (**Supplementary Fig. 1A**). And permutation testing further confirms scale-dependent organization of neuronal diversity (p-value < 0.01; **Supplementary Fig. 1B**). Different brain regions diversify neuronal morphology at distinct hierarchical levels (**Fig.4D**), e.g tract level for AId while arbor level for CP. Cortical neurons exhibit maximal diversification at both primary axonal-tract and full morphology level as most maximal entropy values of all 4 levels S > 0.7 except SSp-ul and SSp-ll, consistent with the need to route signals to diverse targets. Thalamic neurons diversify at full morphology level with all S > 0.75 except LGd at predicted pre-synaptic site level and RT at arbor level, indicating that cell type identity in thalamus is apparently defined by circuit-scale wiring configurations rather than local morphological structures. In contrast, striatal neurons show the largest diversification at the arbor level (S>0.8), suggesting that branch-level organization is a primary axis of morphological specialization in CNU. Based on these, we could ^(^formalize into a simple rule: 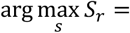 *functional specialization scale of region r*. Different ^&^morphological scales reveal distinct and non-redundant information about neuronal diversity, whereas single-scale morphology is insufficient to capture such diversity (**Supplementary Fig. 1C**).

Collectively, these findings establish MMB as a general framework for linking hierarchical neuronal structure to regional specialization and circuit identity, while uncovering region-dependent principles of morphological diversification. By providing a multiscale, interpretable representation, MMB offers a direct mechanistic bridge between neuronal structure, circuit organization, and functional specialization across the brain.

### MMB reveals a symbolic, quantitative mapping from neuronal structures to brain functions

Neuronal morphology is tightly related to circuit function, and neuronal function is largely reflected in their projection targets. Therefore, we examined how multiscale morpho-barcodes relate to projection across the brain. To this end, we compared correlation matrices derived from MMB pattern distributions and from projection strength profiles across brain regions.

MMB-based clustering clearly segregates brain into major anatomical divisions, aligned with CTX, CNU, and TH, as observed in **Fig. 4A**. However, similarity analyses based on projection strength distributions fail to distinguish comparably distinct anatomical groups (**Fig. 5A**). Correspondingly, Mantel tests reveal weak correlations between morphology-based and projection-based similarity matrices either across the whole brain or for major anatomical groups (CTX, TH, CNU), showing that MMB provides stronger discriminatory power for anatomical organization than projection strength alone. Yet cortical neurons display relatively strong correspondence between morphological similarity and projection similarity, showing tighter alignment between structure and output routing in cortical circuits than subcortical systems. These results uncover that MMB captures structural principals partially explained by projection topology, highlighting MMB as a complementary framework for brain-wide circuit classification.

**Fig 5.**
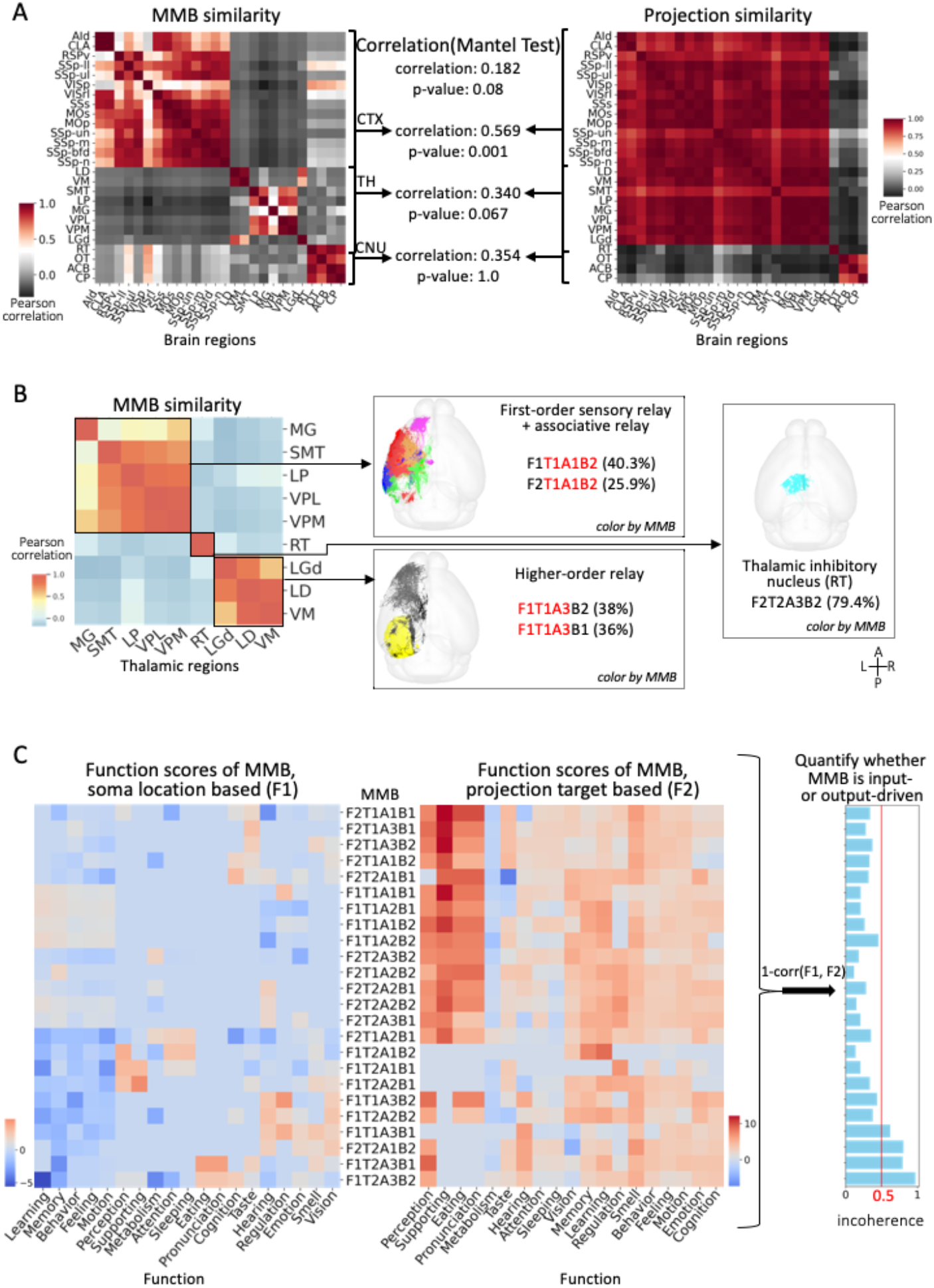
MMB captures anatomical organization beyond projection patterns and functional associations. **A**. Left, similarity matrix across brain regions based on MMB composition. Middle, Mantel tests quantifying correlation between MMB-based and projection-strength-based similarity matrices across all 24 major brain regions, and within CTX, TH and CNU. Right: Similarity matrix based on regional projection strength profiles. **B**. The heatmap shows morpho-pattern similarity among major thalamic regions. Three clusters corresponding to canonical thalamic functional classes identified: i) first-order sensory or associative relay nuclei, ii) higher-order relay nuclei, and iii) inhibitory thalamic nuclei. **C**. Association between morpho-barcodes and brain functions. Heatmaps display correlations between MMB and brain functions denoted as functional scores, based on soma locations (left, correlation denoted as F1) as well as target regions (right, correlation denoted as F2). The bar plot summarizes an incoherence where incoherence = 1-Corr(F, F2) (**Method**), integrating both correlations; a threshold of 0.5 distinguish morpho-patterns predominantly shaped by local (input-driven) versus target (output-drive) constraints.

To further validate the discriminative power of morpho-barcodes, we clustered thalamic neurons based on the similarity of their MMBs. This analysis revealed 3 functionally relevant classes: first-order relay (TH1), higher-order relay (TH2) and inhibitory nucleus (TH3), consistent with canonical thalamic circuit divisions [32] (**Fig. 5B**). These classes exhibit distinct barcode associations: TH1 (SMT, LP, MG, VPL, VPM) maps to F1T1A1B2, F2T1A1B2 modules; TH2 (LGd, LD, VM) aligns with F1T1A3B2, F1T1A3B1 modules; and TH3 (Reticular nucleus of the thalamus, RT) corresponds to F2T2A3B2 module. Thus, the symbolic MMB representation accurately recapitulates functionally defined thalamic organization, demonstrating that multiscale morphology encodes sufficient information to reveal functional circuit hierarchies especially within thalamic regions.

Finally, we sought to quantify how morpho-barcode relate to neuronal functional identity [33] by constructing a morphology-function association mapping. Functional labels were assigned using regional functional annotations from prior biological knowledge. Two complementary maps were generated: one based on soma locations and the other based on projection target regions (**Fig. 5C**). By introducing a ratio metric integrating both maps, we derived an incoherence score that quantifies whether morpho-barcodes are more strongly associated with neuronal input location or output targets. Only four barcodes, i.e. F1T1A1B1, F1T1A1B2, F2T1A1B1, F2T1A1B2, have scores exceeding 0.5, indicating that these four patterns are predominantly driven by input requirements rather than downstream connectivity. Conceptually, this relationship can be framed as an input-output constraint principle: *Morphology* = *α · input constraint* + (1 − *α*) · *output constraint,where α* ≈ *I* (*incoherencedefined above*). This formulation provides a quantitative interpretation of how structure balances afferent and efferent demands of neurons, and may be further generalized and validated with larger datasets. Neurons of these four MMBs correspond primarily to first-order sensory and associative relay nuclei of the thalamus, including SMT, LP, VPM, VPL, and MG [32]. Morphologically, they are characterized by long primary axonal tracts as well as complex proximal axonal arbors and long-distance distal axonal arbors. Such structures without apical dendrites are optimized for receiving specific afferents instead of hierarchical cortical integration. Collectively, T1A1 motif is identified as a conserved thalamic morphotype shared across multiple sensory nuclei, revealing a common structure underlying first-order thalamic relay function.

## DISCUSSION

Neuronal morphology is fundamental to circuit function, yet its complexity has limited systematic, brain-wide analysis. In this study, we introduce multiscale, symbolic morpho-barcoding, a unified framework for quantifying and interpreting neuronal morphology across hierarchical scales at the whole-brain level. Applying MMB to a standardized dataset of whole-brain neuronal reconstructions confirms that neuronal morphology follows hierarchical and region-specific organizational principles [20], and reveals robust associations between morphology, projection patterns, and functional specialization (**Figs. 4–5**).

A key advance of our work is the integration of subcellular and macroscale morphometric features within a single representation, spanning full neuronal morphology as well as the primary axonal tract, arborization, and predicted presynaptic sites (**Fig. 2**). Previous studies have examined modularity at the macroscale using magnetic resonance imaging [34], or at the mesoscale through neuronal population classifications based on projection networks [5, 35, 36, 37]. MMB extends these efforts to the subcellular level. By combining codes across four distinct scales, MMB captures both conserved structural motifs and morphological diversity across brain regions (**Figs. 3–4**), while enabling direct interpretation against biological annotations. For example, arbor complexity correlates with soma location and molecular identity; primary axonal tract structure aligns with long-range connectivity; and predicted presynaptic site distributions most strongly reflect projection type.

By discretizing morphology into combinatorial morpho-patterns across scales, MMB provides a compact yet expressive vocabulary for neuronal structure. We identify 24 unique multiscale morpho-patterns (F#T#A#B#), each representing a specific combination of structural motifs (**Fig. 2B; Fig. 3A**). These morpho-patterns show pronounced regional biases. Thalamic neurons exhibit greater heterogeneity and functional subdivision than cortical and striatal neurons (**Fig. 3**) [38, 39]. Cortical neurons diversify primarily at the level of primary axonal tracts, consistent with the cortex’s role in distributed information integration. Striatal neurons show maximal variability in arbor organization. In contrast, thalamic neurons diversify across both primary axonal tract features and predicted presynaptic-site features, highlighting their diverse relay and modulatory roles. Together, these results indicate that morphological diversity emerges preferentially at specific structural scales, contributing to functional specialization.

Our findings also support and refine prior observations. For instance, reticular thalamic (RT) neurons are known to mediate thalamocortical inhibition and implicated in absence epilepsy [40, 41]. They are predominantly the A3 arbor type, distinguishing them from other thalamic principal neurons and potentially relating to their distinctive roles in sleep and attention [42]. Similarly, the three major morpho-patterns identified here (**Fig. 5B**) correspond closely to classical thalamic classes, including bushy neurons, radiate neurons, and interneurons [42], providing a quantitative bridge to established taxonomy.

Beyond projection topology, MMB captures additional structural information critical for anatomical and functional discrimination. Similarity analyses show that MMB separates major anatomical divisions more robustly than projection-strength profiles alone, and that the correspondence between morphology and projection varies across brain regions (**Fig. 5A**). By integrating soma-based and projection-based functional maps, MMB further demonstrates that distinct morpho-barcodes are shaped by different combinations of local and long-range constraints: some appear predominantly input-driven, whereas others align more strongly with output targeting. Thus, neuronal morphology reflects an interplay between anatomical context and circuit function.

Furthermore, we compared MMB with previously defined c-types [39] and an atlas derived from dendritic microenvironment (ME) [38], to evaluate the its role in resolving neuronal identity and biological significance. Taking VPM neurons as an instance (**Supplementary Fig. 2**), no one-to-one or many-to-one correspondences were observed between MMB and c-type as well as ME based classifications. Instead, two c-types shared similar MMB compositions, and neurons of F1T1A1B2 module were distributed across all VPM ME-subregions. Mutual information between MMB and either c-types or ME defined subregions remained low (<0.5; **Supplementary Table 1**), indicating that MMB captures an independent yet biologically meaningful axis of variation such as associations with functional motifs mentioned above (**Fig. 5C**). This supports the view that neuronal organization is intrinsically multimodal, emerging from the combined constraints of transcriptomic identity, connectivity and anatomy, rather than any single modality alone. Through encoding multiscale morphological structure into a symbolic representation, MMB enables direct computational comparison and integration across modalities, thus offering a novel representation and characterization way for multimodal neuroinformatics. In addition, limitations in imaging precision and spatial registration [45, 46, 47] introduce deviations in localization, preventing reliable assignment of some neurons to the ME atlas. Thus, MMB offers a complementary and robust strategy for characterizing neuronal structure independent of spatial information. In the subsequent research, we will conduct a more comprehensive analysis to illustrate this point.

Together, our results establish MMB as a general framework for studying structure–function relationships across the brain. By transforming complex morphologies into interpretable symbolic representations, MMB enables scalable comparisons across regions and suggests how hierarchical morphological patterns balance structural stereotypy with functional diversity. Ultimately, MMB offers a new approach for uncovering principles of neuronal organization, clarifying links between neuronal structure and function, and advancing our understanding of how the brain works.

## CODE AVAILABILITY

The analytical source codes are available at https://github.com/SEU-ALLEN-codebase/MMB. Dependencies are listed in the requirement.txt and can be easily installed using pip. Detailed documentation, including step-by-step instructions, is provided in the repository.

## DATA AVAILABILITY

All morphometry data and the two **Tables 1 and 2** are accessible from Google Drive at https://drive.google.com/drive/folders/1aF5njPqDW9v-TX2kQdJ3k5-KzSJ2JyeN.

## ACKNOWLEDGEMENT

This work was mainly supported by several grants awarded to H.P., who is a New Cornerstone Investigator and a Shanghai Academy of Natural Sciences (SANS) Senior Investigator. We thank members in Peng Lab for contribution of early development and discussion of this work.

## AUTHOR CONTRIBUTIONS

H.P. conceptualized and managed this study. S.Z. performed data analyses with the assistance of Y.L. Y.L. helped supervising this study and contributed to method development. S.Z. wrote the initial manuscript, which was intensively discussed and rewritten by coauthors.

## COMPETING INTERESTS

The authors declare no competing interests.

## METHOD

### Nomenclature

The nomenclature of brain regions follows CCFv3 [28], which divides a mouse brain into 671 regions. Except for the direct tectospinal pathway (tspd), each region compromises two mirrored subregions in the left and right hemispheres. A finer granularity comprises 314 regions (CCF-R314), achieved by merging highly homogeneous regions. All brain regions used in this paper are derived from the CCF-R314 regions. The CCFv3 atlas is available at https://connectivity.brain-map.org/3d-viewer?v=1. Supra-regional anatomical entities, such as brain areas, are collections of spatially contiguous and functionally related areas. In this paper, we discussed a higher granularity involving 4 functional groups: cortex (CTX), cerebellum (CB), cerebral nuclei (CNU), and brain stem (BS). For CB, we focused on the compound area thalamus (TH), considering our dataset.

### Data acquisition and processing

We used the dataset consisting 1876 single-neuron morphologies (SEU-A1876) [16] manually annotated and corrected based on the CAR platform [26], mapped onto 25um CCFv3 template with the tool mBrainAligner [29, 30]. Each reconstruction follows a strict quality control procedure involving manual cross-validation, automatic quality inspection, and skeleton refining.

Subcellular structures were extracted accordingly from full morphologies to enable a multiscale structural analysis. The primary axonal tract was identified by iteratively removing branches shorter than the second longest branch along the longest axonal path. Axonal arbors were detected using the tool AutoArbor, which employs spectral clustering to identify densely packed sub-tree structures within the full morphology. To ensure comparability, the dominant number of arbors for neurons within the same brain region was pre-determined using a majority-vote approach. Varicosity detection was performed in two main steps: determining initial candidates which show overlapped peaks in the intensity and radius profiles of axonal skeleton against neuronal images; filtering out false positive results with radius no more than 1.5 times of its surrounding axonal nodes [43, 44] or intensity value lower than 120 in 8-bit images (maximum intensity 255), and possible duplicates within five voxels in highest-resolution, roughly equivalent to the size of a typical varicosity (1-2um).

### Vaa3D based L-Measure features

We used L-measure [21] features to characterize full morphology. We computed 22 features from the “global_neuron_feature” plugin in Vaa3D, including: “Nodes,” “SomaSurface,” “Stems,” “Bifurcations,” “Branches,” “Tips,” “OverallWidth,” “OverallHeight,” “OverallDepth,” “AverageDiameter,” “Length,” “Surface,” “Volume,” “MaxEuclideanDistance,” “MaxPathDistance,” “MaxBranchOrder,” “AverageContraction,” “AverageFragmentation,” “AverageParent-DaughterRatio,” “AverageBifurcationAngleLocal,” “AverageBifurcationAngleRemote,” and “HausdorffDimension.”. While these L-Measure Vaa3D features are conceptually similar to those defined in the L-Measure server (http://cng.gmu.edu:8080/Lm), some differences exist in their implementation.

### Morphological feature extraction and quantification

We computed 22 features for full morphologies utilizing Vaa3D plugin “global_neuron_feature”, excluding five inaccessible features (e.g., soma surface and total surface area) for manually annotated neurons. This resulted in a 47-dimensional feature vector, comprising 7 global features and 10 local features, where each local feature was quantified using four statistical metrics: minimum, maximum, mean and standard deviation.

Morphological features for subcellular structures:

1. 6 primary axonal tract features: total path length, Euclidean distance to soma, orientation length along 3 axis and total volume.
2. 9 features for each arbor structure: number of nodes (“#node”), number of branches (“#branch”), total arbor volume (“volume”), total path length, distance to soma, number of hubs (“#hub”), and maximal spatial density (“max_density”) defined as the number of axonal nodes within a 20 µm radius for each node. Based on the Euclidean distance from the maximal density node to the soma, axonal arbors could be classified as axonal arbor 1, axonal arbor 2, …, etc., from near to far.
3. 7 features for predicted pre-synaptic site: number of predicted pre-synaptic sites, ratio of terminaux (TEB) ratio, density of predicted pre-synaptic sites within 1 mm^3^, average Euclidean distance to soma, inter-distance between predicted pre-synaptic sites, number of projecting regions, number of segments.

All features were standardized using Z-score normalization. Neuronal similarity was evaluated based on Euclidean distances between these feature vectors.

### MMB generation

Based on the feature matrix of each morphological level, we performed Hierarchical Clustering and determined the optimal number of clusters according to Silhouette scores. This yielded 2 clusters at full morphology level, 2 at primary axonal tract level, 3 at arbor level, and 2 at predicted pre-synaptic site level. Combining these four levels, we obtained the barcode represented in the format ‘F#T#A#B#’, where # denotes the cluster ID for the corresponding level.

### Functional score

The functions of each brain region here are listed by biologists. For each Multiscale-morpho-barcode *m* and brain function *f*:

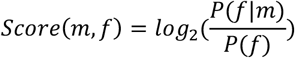

Where:

P(*f*|*m*): fraction of neurons of MMB *m* in regions associated with *f*.

P(*f*): overall fraction of neurons associated with f across all neurons.

For each Multiscale-morpho-barcode m and its projection t:

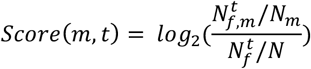

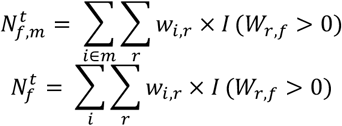

Where:

*W*_*r,f*_ ∈ [0,1], 1 if region r associated with function f.

*w*_*i,r*_: projection strength ratio as normalized weight.

*N*_*m*_: number of Multiscale-morpho-barcode m across all neurons.

*N*: total number of neurons

## Notes

### Competing Interest Statement

The authors have declared no competing interest.

