## Supplementary figures and images for "Multiscale Symbolic Morpho-Barcoding Reveals Region-Specific and Scale-Dependent Neuronal Organization"

### Supplementary Fig.1

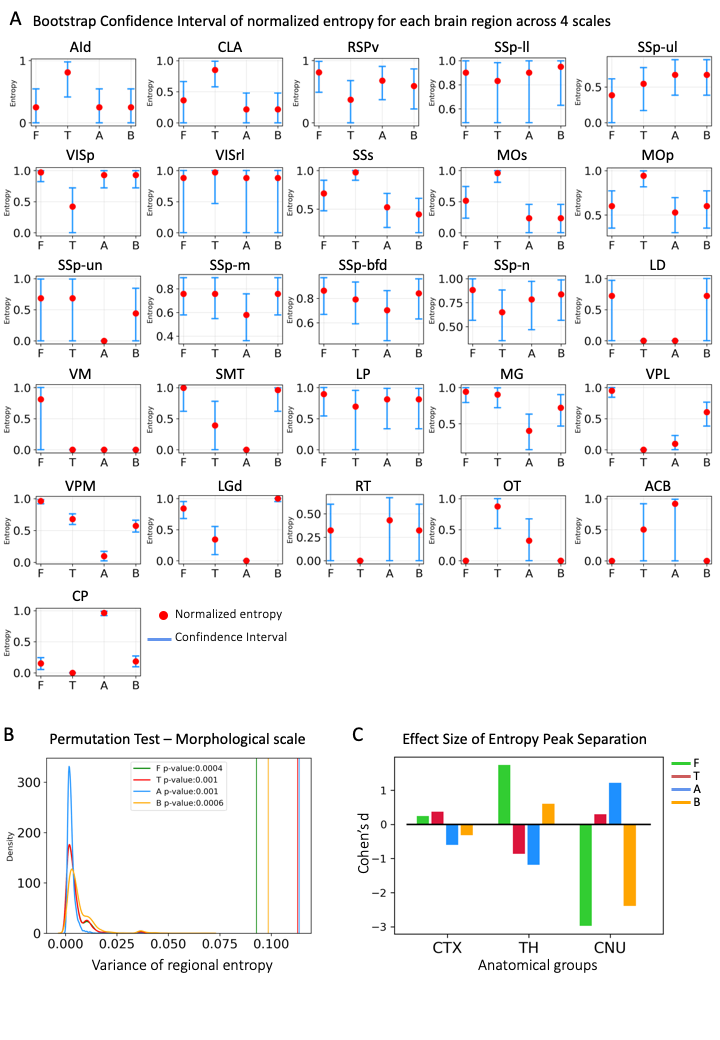

### Supplementary Fig.2

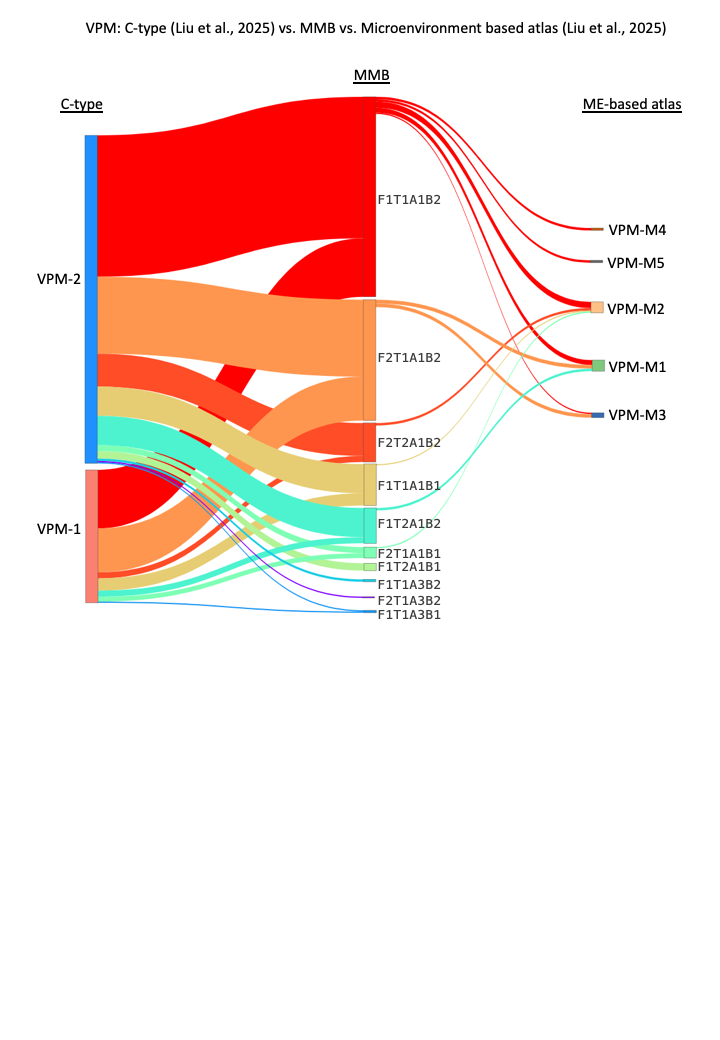
